# Aerobic Bacteria Produce Nitric Oxide via Denitrification and Trigger Algal Population Collapse

**DOI:** 10.1101/2021.11.14.468512

**Authors:** Adi Abada, Martin Sperfeld, Raanan Carmieli, Shifra Ben-Dor, Irene Huang Zhang, Andrew R. Babbin, Einat Segev

## Abstract

Microbial interactions govern marine biogeochemistry. These interactions are generally considered to rely on exchange of organic molecules. Here we report on a novel inorganic route of microbial communication, showing that algal-bacterial interactions are mediated through inorganic nitrogen exchange. Under oxygen-rich conditions, aerobic bacteria reduce algal-secreted nitrite to nitric oxide (NO) through denitrification, a well-studied anaerobic respiration mechanism. Bacteria secrete NO, triggering a cascade in algae akin to programmed cell death. During death, algae further generate NO, thereby propagating the signal in the algal population. Eventually, the algal population collapses, similar to the sudden demise of oceanic algal blooms. Our study suggests that the exchange of denitrification intermediates, particularly in oxygenated environments, is an overlooked yet ecologically significant route of microbial communication within and across kingdoms.

**One Sentence Summary:** Aerobic bacteria activate denitrification in oxygenated conditions and produce nitric oxide that kills their algal partners

## Main text

Various marine bacteria that inhabit oxygenated surface waters carry denitrification genes. This is surprising, as denitrification is a microbial process which allows microorganisms to maintain cellular bioenergetics in limiting oxygen concentrations (*1*). To conduct denitrification, specialized microbes express enzymes that are responsible for a multi-step reduction process of nitrogen species that serve as terminal electron acceptors. The full denitrification process commences with the reduction of nitrate (NO_3_^−^) to nitrite (NO_2_^−^), then to nitric oxide (NO), to nitrous oxide (N_2_O) and finally to dinitrogen (N_2_) (Fig. 1A). In the ocean, microbial denitrification widely occurs in oxygen deficient zones (ODZs) and ocean sediments (*2, 3*), and all denitrification intermediates including nitrite, NO and nitrous oxide, are observed to accumulate in the marine environment (*3-5*). Knowledge regarding the NO intermediate is scarce, however, due to its short-lived nature (*3*). Therefore, its cryptic presence in the ocean water might have overlooked consequences.

**Fig. 1.**
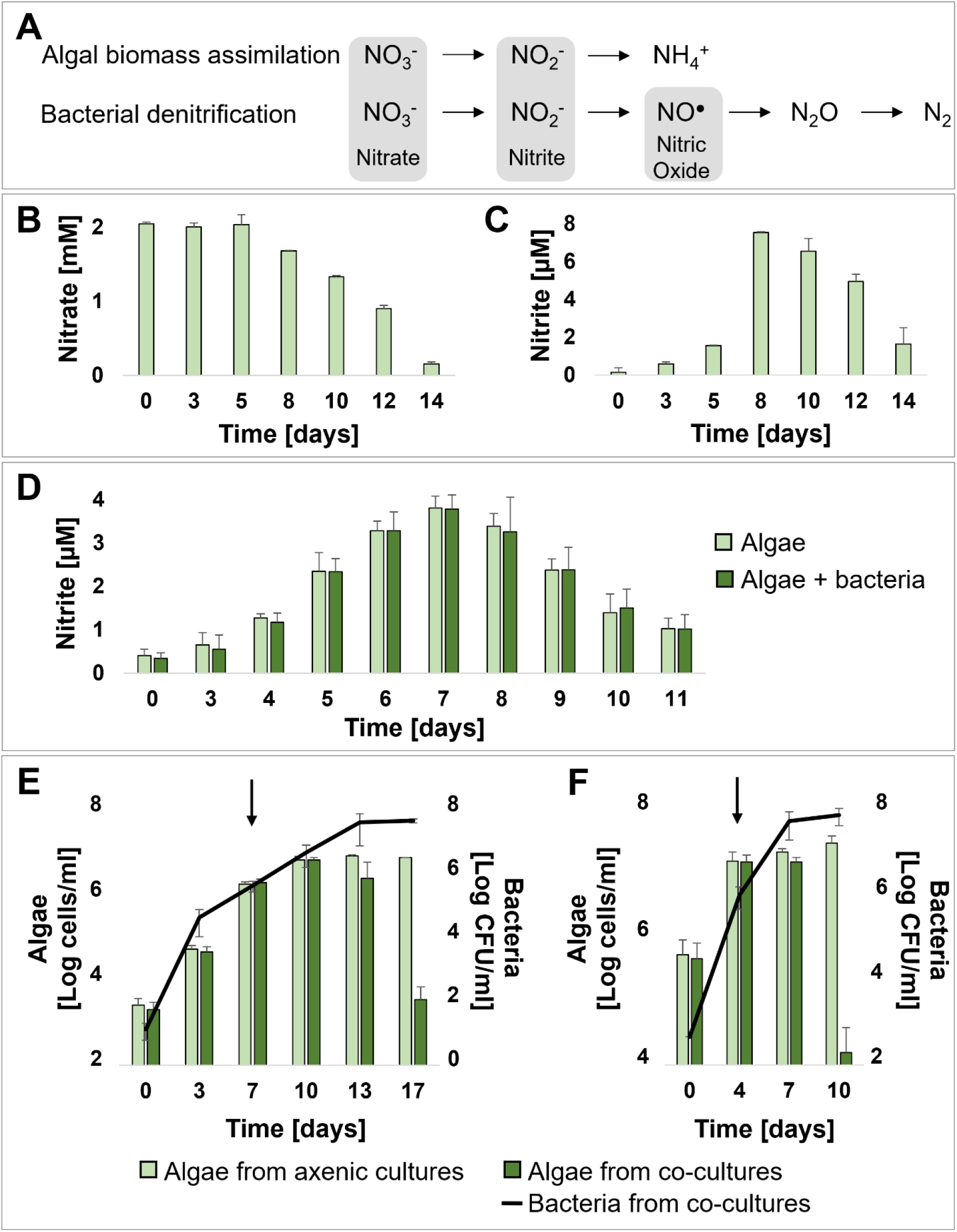
Algal nitrite secretion during exponential growth is linked to the timing of algal death in co-cultures. **(A)** Assimilatory nitrate reduction to ammonium during algal biomass assimilation produces nitrite as an intermediate. Dissimilatory nitrate reduction to di-nitrogen during bacterial denitrification also includes nitrite as an intermediate, which is further reduced to NO. The main nitrogen species discussed in the manuscript are shaded in gray. (**B**) Nitrate and (**C**) nitrite detected in filtrates of axenic algal cultures on the indicated days. (**D**) Daily monitoring of nitrite in filtrates of axenic algal cultures (light green) and co-cultures (dark green). **(E, F)** Algal growth (bars) and bacterial growth (black line) in axenic algal cultures (light green bars) and algal-bacterial co-cultures (dark green bars). (E) Cultures initiated with an algal inoculum of 10^3^ algae/ml. (F) Cultures initiated with an algal inoculum of 10^5^ cells/ml. Black arrows indicate the day of the extracellular nitrite peak as shown in (D) and (S1D). Each data point in the figure consists of at least 3 biological replicates, error bars designate ± SD.

NO in the ocean has origins in addition to denitrification. In fact, most living organisms can produce this molecule for signaling, both under normal and perturbed physiology (*6-8*). NO is often associated with processes of programmed cell death (PCD) and oxidative stress, both in unicellular and multicellular organisms (*9-12*). Since NO is a small, membrane-permeable gas molecule, it passes easily through adjacent cells (*13*). The short half-life of NO in oxygenated environments ensures a localized response within the tissue (*13*). Similarly, NO in unicellular organisms can stimulate a response in a close neighbor (*14-16*). Indeed, NO has been suggested to diffuse between cells in a dense population of the microalga *Emiliania huxleyi* (*17*).

*E. huxleyi* is the most widespread coccolithophore in modern oceans (*18*). Coccolithophores are unicellular marine algae that cover their cell with intricate platelets made of crystalline calcium carbonate. *E. huxleyi* forms vast annual blooms that can stretch over thousands of square kilometers of ocean surface (*19, 20*). The blooms gradually form over several weeks and then suddenly collapse (*21-23*). NO has been shown to play a role in bloom demise, particularly as a result of viral infection (*17*). Notably, algal populations that exhibit NO production in the ocean and in the lab also harbor a rich bacterial community. Among these bacteria are the Roseobacters, an abundant and metabolically versatile group of marine bacteria that are commonly found associated with microalgae (*24-28*). Many Roseobacter bacteria carry denitrification genes (*29*), and thus have the potential to generate NO and contribute to the observed NO-related algal demise. However, many members of this bacterial group are strict aerobes (table S1), thus the presence of denitrification genes in their genome is puzzling.

Here, we use an algal-bacterial model system, comprised of the alga *E. huxleyi* and the Roseobacter bacterium *Phaeobacter inhibens* (*28*), to study microbial inorganic nitrogen exchange. The interaction between *E. huxleyi* and *P. inhibens* is dynamic; initially algae and bacteria exchange beneficial molecules in a mutualistic phase (*28*). However, as the cultures age, bacteria become pathogenic and kill their algal partners (*28*). We show that the strict aerobic Roseobacter bacterium *P. inhibens* produces NO through denitrification under well-oxygenated conditions, similar to the global surface ocean. Our data reveal that *P. inhibens* bacteria generate NO by reducing nitrite that is secreted by the alga *E. huxleyi* during algal exponential growth. Furthermore, we demonstrate that bacterial NO is detected extracellularly, and triggers a PCD-like process in the algal population. As part of their death process, algae further generate and release NO, thereby propagating the signal among the algal population. Importantly, environmental metagenomic data and analysis of metagenome-assembled genomes (MAGs) from oxygen-rich waters support the co-occurrence of bacteria that carry denitrification genes and phytoplankton, highlighting the potential ecological relevance of our laboratory findings. Our results unveil an algal-bacterial chemical crosstalk mediated through inorganic nitrogen species and point to NO as an inter-kingdom signaling molecule. These observations have implications for many interactions between eukaryotic hosts and bacteria that harbor denitrification genes for purposes other than anaerobic energy production.

## Results

### Algae secrete nitrite in a growth phase-dependent manner

Algal exudates of *E. huxleyi* are necessary to enable growth of the bacterium *P. inhibens* in seawater *(28)*. While it is commonly known that algal nutrients can support the growth of heterotrophic bacteria (*30*), the identity and quantity of inorganic nutrients that are secreted by specific algae, remain unknown. To better understand the algal-bacterial interaction, we measured the main inorganic nutrients that are secreted by growing algal cultures. Our data indicate that the levels of nitrate (NO_3_^−^) and phosphate (PO_4_^3-^) in the medium gradually decreased over time (Fig. 1B, fig. S1A), indicative of assimilation by the growing algal population. As algae are primary producers, they can uptake inorganic nitrogen from seawater in the form of nitrate, reduce it to nitrite (NO_2_^−^) and further reduce it in an assimilatory pathway to generate ammonium (NH_4_^+^) (Fig. 1A). Surprisingly, we observed an extracellular peak of the intermediate, nitrite (Fig. 1C). Nitrite was previously reported to leak from diatoms, another microalgal group, during exponential growth. The exuded nitrite can later be up-taken by cells at stationary phase (*31-33*). Contrary to the nitrite peak, ammonium and sulfate (SO_4_^2-^) levels remained unchanged in the medium (fig. S1B-C).

To gain a greater temporal resolution of algal nitrite secretion, we monitored daily the extracellular levels of nitrite in axenic algal cultures. The highest nitrite concentration was repeatedly detected during the algal exponential growth phase (Fig. 1D, E). To further characterize the association between algal growth and nitrite dynamics, we initiated algal cultures with a high algal inoculum (10^5^ cells/ml) (Fig. 1F, fig. S1D). A denser algal inoculum results in expedited exponential growth and an earlier stationary phase (Fig. 1F). Consequently, in cultures initiated with a denser inoculum, the nitrite peak was detected earlier than in cultures that were initiated with a more dilute inoculum (fig. S1D). Taken together, these observations suggest that nitrite secretion is indicative of an exponentially growing algal population.

Bacteria substantially impact algal physiology and influence growth dynamics of algal populations (*28, 34-36*). We therefore explored whether co-culturing algae with bacteria would influence nitrite secretion. The concentration and timing of the nitrite peak was notably similar in axenic algal cultures and algal-bacterial co-cultures, regardless of the initial algal inoculum (Fig. 1D, fig. S1D). These findings suggest that nitrite secretion by algae is not influenced by the presence of bacteria, and remains an indicator of algal growth phase in co-cultures.

### Algal death in algal-bacterial co-cultures is correlated to the algal nitrite peak

Bacterial pathogenicity is triggered as algal cultures age (Fig. 1E) (*28*). Therefore, we examined whether algal-secreted nitrite can serve as a signal for bacteria to become pathogens (manifested by algal death). In line with this idea, changing the timing of the nitrite peak is expected to result in a concomitant shift in the timing of sudden algal death. Timing of nitrite secretion can be altered by adjusting the density of the algal inoculum (fig. S1D), therefore we initiated algal-bacterial co-cultures with a denser algal inoculum. As can be seen in Fig. 1F, a denser algal inoculum resulted in earlier algal death. These results propose a correlation between the growth phase of the algal population, indicated by the peak of algal nitrite, and the subsequent bacterially-triggered algal death. Taken together, algal nitrite is likely a signaling molecule for bacteria that indicates algal growth phase and triggers bacterial pathogenicity.

### Bacteria harbor denitrification genes related to nitrite metabolism

To explore whether secreted algal nitrite serves as a signaling molecule for bacteria, we investigated the genetic potential of *P. inhibens* to detect and metabolize nitrite. A search for bacterial functions related to nitrite metabolism identified four genes in the *P. inhibens* genome (fig. S2A). Two are nitrite reductases; *nirK*, which has been extensively studied in nitrite reduction to NO in denitrifying bacteria (*37*), and a putative nitrite/sulfite reductase, mainly characterized in sulfur metabolism. The two other detected genes are a nitrite transporter, related to nitrite uptake but not to further metabolism, and a protein containing a nitrate/nitrite sensing domain with no suggested activity. By analyzing the genomic locus of *nirK*, we identified multiple genes and operons that are related to denitrification: an adjacent *nirV* gene, commonly found in the *nirK* operon, yet without known activity (*38*), a *nor* operon encoding an NO reductase (*norCBQD*), a *nnrS* gene that is suggested to help alleviate NO-related stress in bacteria (*39*), and a *nnrR* gene encoding an established NO transcriptional regulator (*40, 41*) (fig. S2B). These denitrification-related functions are all found on a single bacterial 262 kb plasmid, one of three native plasmids of *P. inhibens (42)*.

Denitrification is a common metabolic process in anaerobic bacteria that reduce inorganic nitrogen species as terminal electron acceptors for respiration (Fig. 1A) (*1*). *P. inhibens* specializes in interactions with photosynthesizing hosts that produce oxygen (*28*), therefore it is curious why this bacterium maintains denitrification genes. To address this question, we examined the ability of *P. inhibens* bacteria to grow under oxygen-depleted conditions by utilizing nitrate or nitrite as terminal electron acceptors (Fig. 1A). Our results demonstrated that *P. inhibens* was unable to grow under anaerobic conditions (fig. S2C), while the known facultative anaerobic bacterium *Escherichia coli* did. This result indicates that *P. inhibens* is a strict aerobe that maintains denitrification genes for reasons other than bioenergetics.

### Bacteria can produce and secrete nitric oxide under aerobic conditions

*E. huxleyi* cells secrete nitrite during exponential growth (Fig. 1) and *P. inhibens* bacteria have the genetic potential to reduce nitrite to NO through denitrification (fig. S2A,B). NO was previously shown to play key roles in algal death following viral infection (*17*) and in PCD in multicellular organisms (*6-8*). We therefore explored whether NO is involved in algal death during the *E. huxleyi-P. inhibens* interaction. First, we examined whether denitrification genes can be expressed in pure bacterial cultures under aerobic conditions. Our data show that immediately following exposure to exogenous nitrite, mimicking the nitrite secreted by algae, the expression of bacterial *nirK* and *norB* increased, reaching an expression peak of 4-fold and 10-fold, respectively, within one hour of exposure (Fig. 2 A, B). The enzyme encoded by *nirK* produces NO from nitrite while the enzyme encoded by *norB* further reduces NO within a bacterial cell (Fig. 1A).

**Fig. 2.**
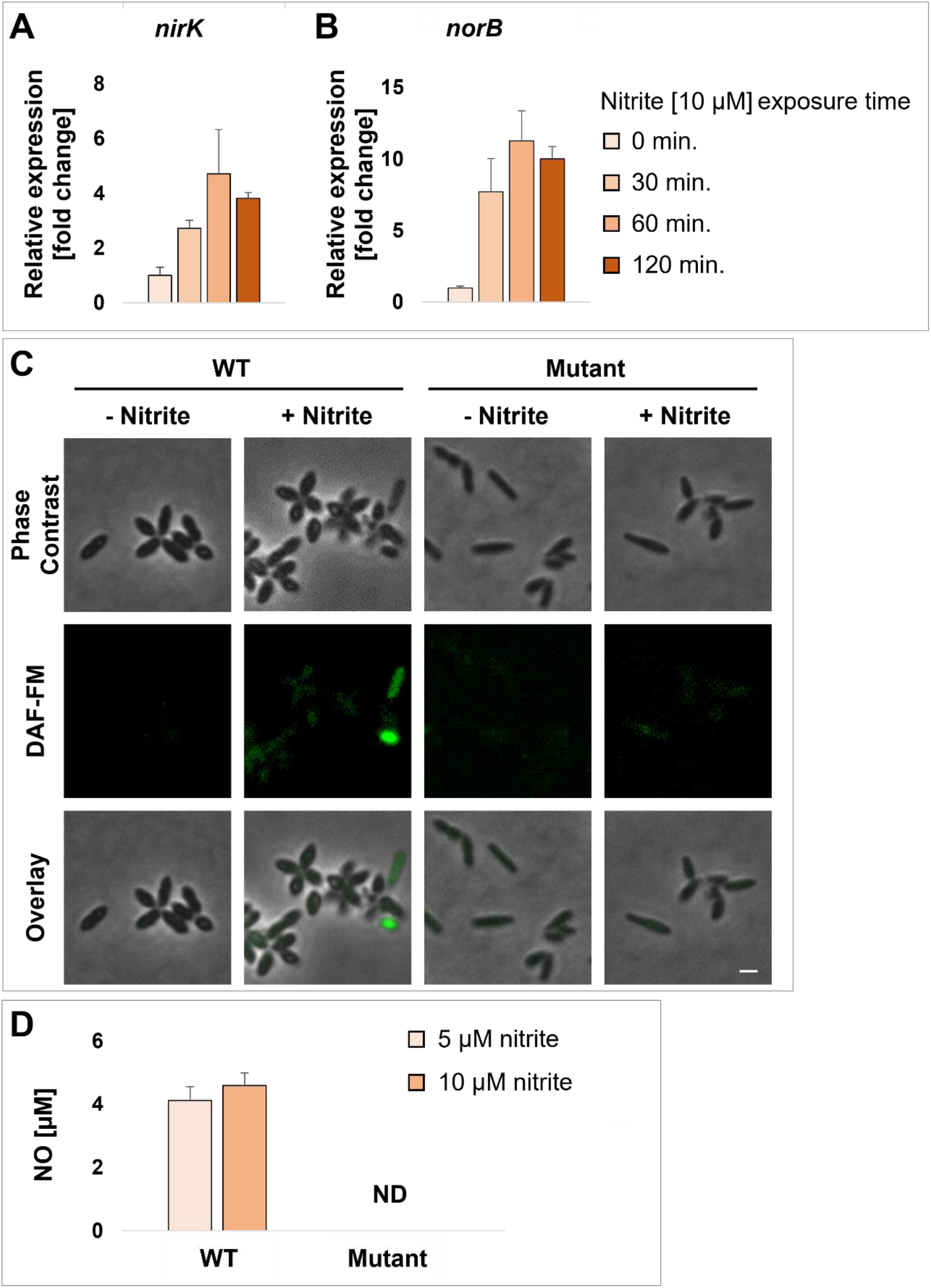
Bacteria produce and secrete NO under aerobic conditions. (**A, B**) Relative gene expression of *P. inhibens* denitrification genes under exposure to 10 µM nitrite for the indicated times; (A) *nirK*, (B) *norB*. Each data point consists of 3 biological replicates, error bars designate ± SD. (**C**) Microscopy images of WT and mutant bacteria cured from the native 262 kb plasmid, stained with the fluorescent NO-indicator diacetate DAF-FM, and incubated with or without 100 µM nitrite for 2 h. Scale bar corresponds to 1 µm. (**D**) Extracellular nitric oxide concentrations of WT and mutant bacteria incubated with the indicated nitrite concentrations. Each data point consists of 3 biological replicates, error bars designate ± SD. ND stands for not detected.

Next, we tested whether in addition to gene expression, we can detect the production of NO in bacterial cells exposed to nitrite, under aerobic conditions. To this end, intracellular NO production in bacterial cells was tracked using DAF-FM Diacetate (4-Amino-5-Methylamino-2’,7’-Difluorofluorescein Diacetate), a fluorescent probe that emits a signal upon intracellular NO binding. Our results demonstrate that the addition of nitrite to pure bacterial cultures, results in increased levels of intracellular NO (Fig. 2C). To validate that increased NO production is dependent on the expression of denitrification genes, we examined a bacterial mutant that was cured of its native 262 kb plasmid, and therefore no longer carries any denitrification genes (*42*). Our results indicate that under the same experimental conditions, the bacterial mutant grows to the same optical density as the wild-type (WT) (fig. S2D) but does not produce NO (Fig. 2C).

If bacterial NO is indeed involved in algal death, it has to diffuse out of the producing bacterial cell to the extracellular environment, where it can potentially reach and affect neighboring algal cells. To explore this idea, we first assessed whether bacterial NO can be detected extracellularly. Therefore, we utilized the Liposome-Encapsulated-Spin-Trap (LEST) method that was previously developed to measure extracellular NO secreted by microorganisms to their surroundings (*43*), and was successfully applied in microalgal cultures (*17, 43*). Extracellular NO was indeed detected in the medium of bacterial cells supplemented with nitrite, but not in the medium of mutant cells cured of the 262 kb plasmid, or WT cells that were not supplemented with nitrite (Fig. 2D). Taken together, our data demonstrate that *P. inhibens* bacteria produce and secrete NO upon exposure to extracellular nitrite under aerobic conditions.

### NO production by bacteria is essential for triggering algal death

*E. huxleyi* algae secrete nitrite during the exponential growth phase (Fig. 1), and the nitrite peak is followed by bacterial-induced algal death. If bacterial denitrification is involved in promoting algal death, the mutant strain should not be able to trigger algal death in co-cultures. Indeed, algae that were co-cultivated with mutant bacteria did not exhibit algal death (Fig. 3A). Importantly, WT and mutant bacteria exhibit similar growth curves both in pure bacterial cultures and co-cultures (Fig. 3A, fig S2D).

**Fig. 3.**
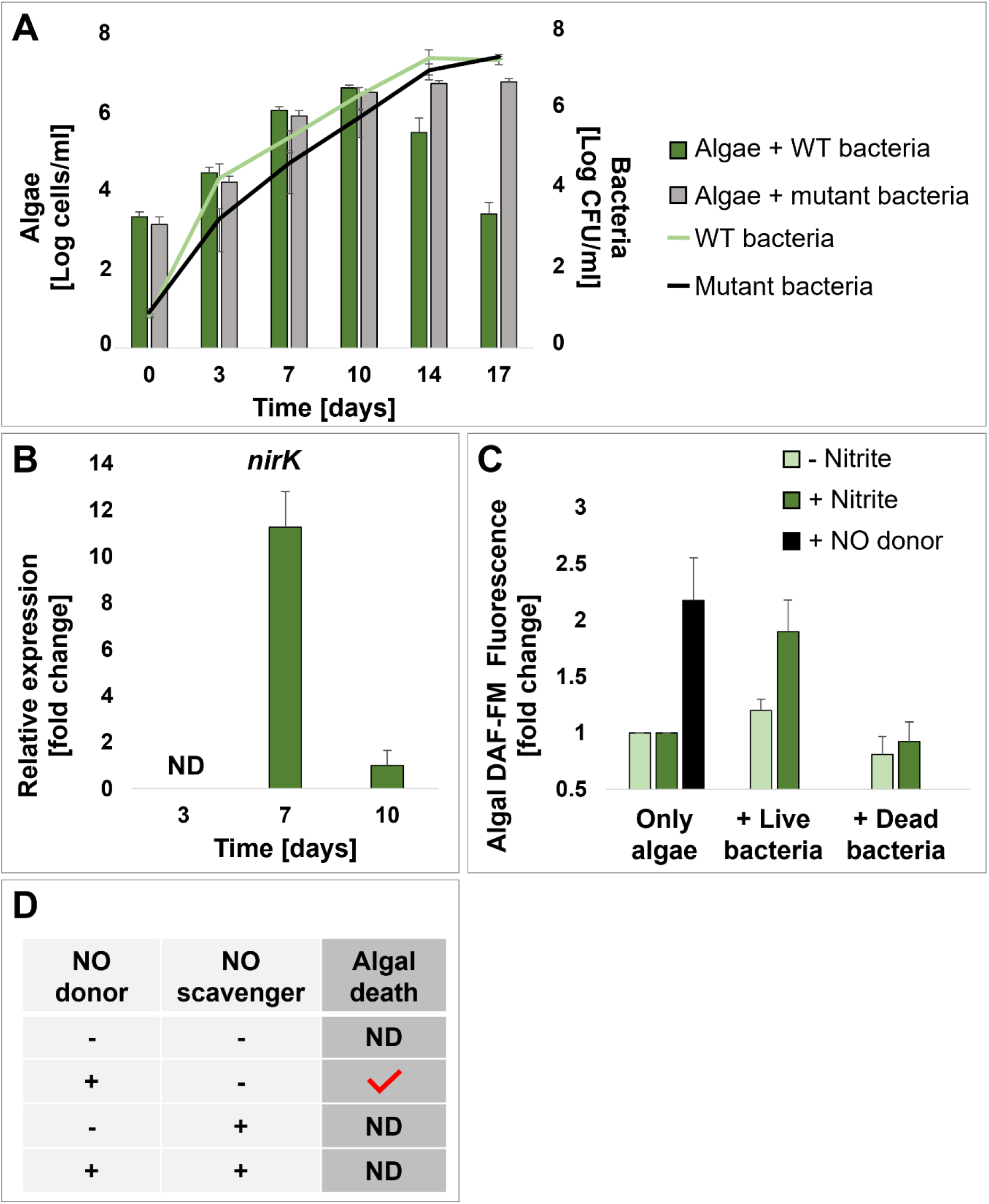
Bacterial NO production by denitrification is essential in triggering algal death. (**A**) Algal growth (bars) and bacterial growth (lines) in co-cultures with WT bacteria (green bar and line) and mutant bacteria cured from the native 262 kb plasmid (gray bar and black line). Each data point consists of 3 biological replicates, error bars designate ± SD. (**B**) Relative expression of *nirK* in bacterial cells from an algal-bacterial co-culture, sampled on the indicated days. Each data point consists of 4 biological replicates, error bars designate ± SD. ND stands for not detected. (**C**) Fluorescence of algal cells stained with the fluorescent NO-indicator diacetate DAF-FM, incubated with live or dead bacteria for 2 h under the following conditions: control (-nitrite, light green), exposure to nitrite (+ nitrite [10 µM], dark green), or exposure to a chemical NO donor (+NO donor, [300 µM] DEANO, black). Results represent 3 biological replicates, each containing 10,000 algal cells, error bars designate ± SD. (**D**) The effect of addition (+) of a NO donor (100 µM DEANO) and a NO scavenger (20 µM c-PTIO) on the death of axenic algal cultures. Red checkmark indicates observed algal death. Results represent at least 3 biological replicates. ND stands for not detected.

To further establish the involvement of bacterial denitrification in triggering algal death, we examined the expression of *nirK*, the gene encoding the NO-producing enzyme in bacteria, in algal-bacterial co-cultures. Expression of this bacterial gene in co-cultures would suggest bacterial NO production through reduction of algal-secreted nitrite. Our results show more than 10-fold increase in *nirK* expression in bacteria in co-cultures during the algal exponential growth phase (Fig. 3B), concurrent with the algal nitrite peak (Fig. 1D, E). We were not able to detect *nirK* expression in co-cultures during earlier culturing stages, either because it is not expressed or due to low bacterial numbers. These results demonstrate the co-occurrence of the algal-secreted nitrite peak, and the response of bacteria that express genes to reduce nitrite to NO in co-cultures.

Bacteria secrete NO when exposed to nitrite (Fig. 2D), and NO is known to diffuse across membranes from producing cells to neighboring cells (*13*). We therefore tested the ability of bacterial NO to diffuse from NO-producing bacteria to adjacent algal cells. We monitored intracellular NO in algal cells that were exposed to DEANO (Diethylammonium (Z)-1-(N,N-diethylamino)diazen-1-ium-1,2-diolate), an extracellular chemical NO donor, or to bacteria. Our results show that when algae were stained with the fluorescent NO indicator DAF-FM diacetate, exposure to an extracellular chemical NO donor resulted in increased fluorescence (Fig. 3C). This observation indicates that external NO can indeed diffuse into algal cells where it binds the fluorescent indicator. Moreover, when algae were incubated with bacteria, and supplemented with nitrite, increased fluorescence was also evident (Fig. 3C). A similar experiment conducted with dead bacteria or without nitrite addition, did not yield increased fluorescence (Fig. 3C). Thus, it appears that live bacteria reduce nitrite and generate NO that can diffuse into adjacent algal cells.

We next examined whether the actual NO molecule triggers algal death. If extracellular NO, secreted by bacteria, causes algal death in the algal-bacterial interaction, treating axenic algal cells with extracellular chemical NO should result in similar sudden algal death. Indeed, addition of a chemical NO donor to axenic algal cultures resulted in death of the cultures (Fig. 3D). This effect was rescued, however, by the addition of the chemical NO scavenger c-PTIO, suggesting that extracellular NO in particular triggers algal death.

### Extracellular NO triggers a specific PCD-like death in algal cells

Previous studies have identified that algae undergo a PCD-like process when algal death is triggered by bacteria (*28, 44*). Several PCD-like and oxidative stress-related genes are upregulated during bacterial-triggered algal death (*28*). If bacterial NO is indeed the inducer of algal death, then extracellular NO should promote upregulation of PCD-related genes. To explore this possibility, we first compiled an updated list of genes that are differentially expressed during bacterial-triggered algal death. Recently, a novel *E. huxleyi* transcriptome was generated by our lab, specifically designed to elicit expression of genes involved in algal-bacterial interactions (*45*). Mining the novel transcriptome revealed various algal genes that are putatively involved in oxidative stress and PCD (fig. S3A). We used these genes as a “fingerprint” for bacterial-induced algal death and monitored changes in the expression of selected genes. Our data show that treatment of axenic algal cultures with a chemical NO donor, mimicking bacterial NO secretion in co-culture, results in increased expression of 3-8 fold of oxidative stress and PCD-related genes which are hallmarks of the bacterially-triggered algal death (Fig. 4A).

**Fig. 4.**
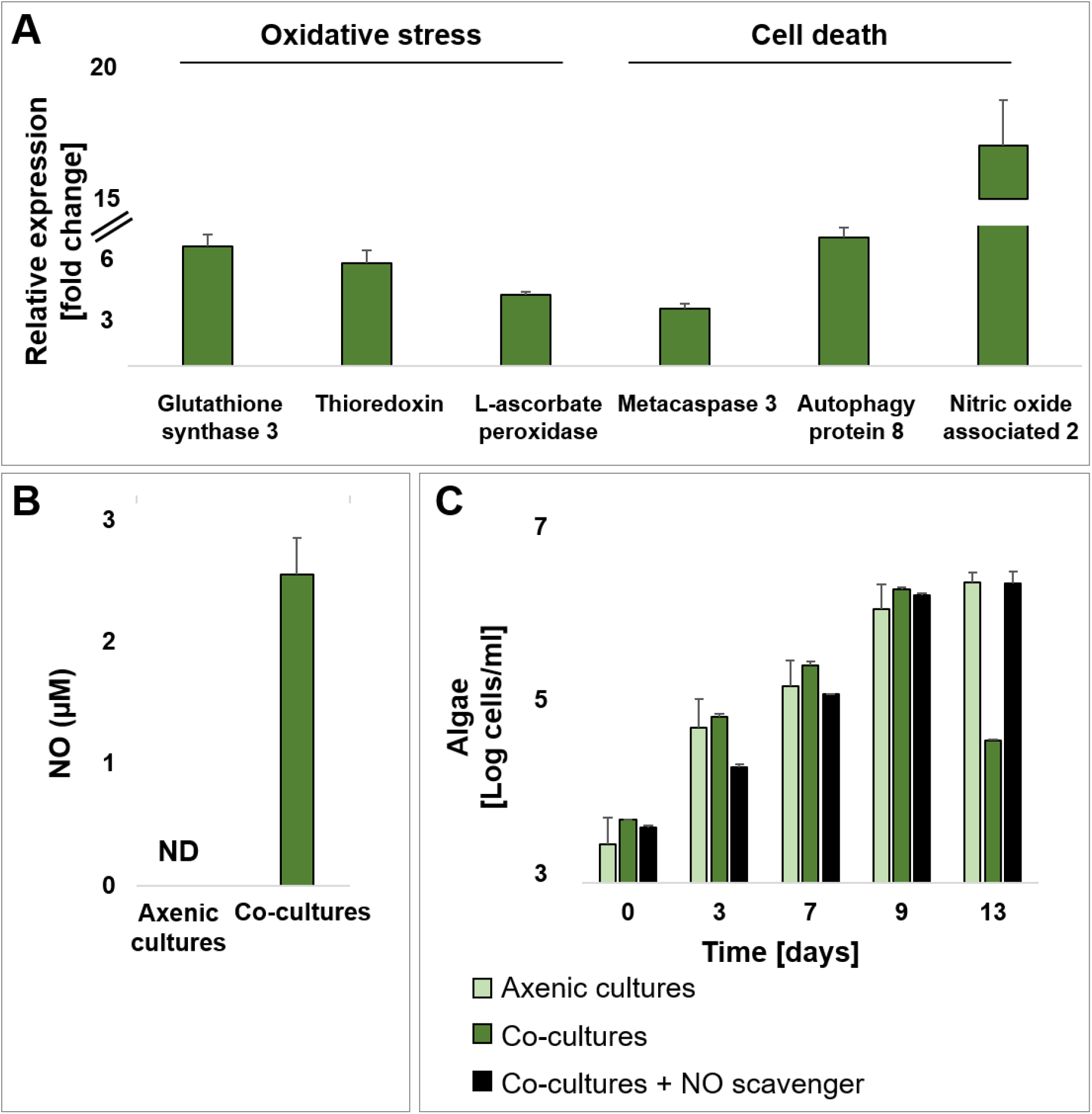
Extracellular NO triggers a PCD-like process in algae, followed by algal NO production and secretion. (**A**) Relative gene expression of algal oxidative stress and PCD-like genes following incubation with a NO donor ([100 µM] DEANO for 18 h). Annotated gene products are depicted. (**B**) Extracellular NO concentrations of axenic algal cultures and algal-bacterial co-cultures on day 10. (**C**) Algal growth in axenic algal cultures (light green), co-cultures (dark green) and co-cultures supplemented with a NO scavenger ([20 µM] c-PTIO added on day 9 of the co-culture, black). All data points in the figure consist of 3 biological replicates, error bars designate ± SD.

### Algae can potentially produce nitric oxide via a newly-identified nitric oxide-associated protein

Algae were shown to produce and secrete NO during cell death following viral infection (*17*). Given that NO appears to be central in the algal-bacterial interaction, we characterized algal NO production, in addition to the bacterial NO source. Despite previous reports regarding the importance of NO production during algal death, the underlying genes were not yet identified. In plants and other microalgae, NO was suggested to be generated by a GTP-binding Nitric Oxide-Associated protein (NOA) (*46, 47*). Using bioinformatics, we now identify two *noa* genes in *E. huxleyi*; *noa1*-similar to the gene sequence of *noa* in diatoms (MZ773649) (*47*) and *noa2*-similar to the *noa* gene sequence in plants (MZ773650) (*46*) (fig. S3B, table S2). Our data indicate that when axenic algal cultures were treated with a chemical NO donor, the expression of the *noa2* gene increased over 15-fold (Fig. 4A). Moreover, as evident in the transcriptome data from co-cultures, the expression of the *noa2* gene increased during stationary phase and algal death, while the expression of *noa1* did not change (fig. S3C) (*45*). Further genetic and biochemical characterization will be needed to underpin the identity and role of these *noa* encoded proteins. Nevertheless, our data show that following exposure to extracellular NO, either from a chemical or a bacterial source, algae upregulate the *noa2* gene that potentially encodes a NO-producing enzyme.

### Algae produce and secrete NO during NO-triggered PCD-like cell death

We further tested whether algae indeed produce NO in algal-bacterial co-cultures. We detected extracellular NO only during the stationary phase of co-cultures, but not in axenic algal cultures of the same age (Fig. 4B). According to our gene expression data, the detected NO is likely of predominant algal origin as the expression levels of the algal *noa2* gene increased during stationary phase in co-cultures (fig. S3C). In light of these findings, it is possible that our previous observation of increased intracellular NO in algal cells exposed to external NO (Fig. 3C), is due to algal NO production. Taken together, our data imply that algal death can be triggered by bacterial NO, and this exogenous NO promotes further NO production and secretion by algae.

Production and secretion of NO by algae can spread the death signal among the algal population; each algal cell that senses external NO becomes a NO-producer, thus amplifying the signal in the population. Hence, algae can propagate the NO death signal to the entire population, resulting in its collapse. A similar mechanism of algal demise was previously suggested to occur following viral infection (*17*). If secreted algal NO indeed spreads as a signal in the algal population, a NO scavenger should rescue algal death in algal-bacterial co-cultures. Indeed, addition of a NO scavenger to co-cultures immediately before the population reaches stationary phase (when the peak of extracellular NO was measured) (Fig. 4B) completely prevented algal death (Fig. 4C). These observations suggest that NO secretion by algae can propagate the death signal among algal cells and promote collapse of the entire algal population.

### Denitrification genes are detected in oxygenated regions of the ocean

The detection and expression of denitrification genes in the ocean is generally associated with denitrifying bacteria in oxygen-depleted regions. However, here we have shown that an obligate aerobic bacterium, *P. inhibens*, expresses denitrification genes while interacting with an oxygenic photosynthesizing host in co-culture. Interestingly, many other Roseobacters are known to be aerobes, often in close interactions with photosynthesizing organisms (*28, 34, 35, 48, 49*), yet numerous strict aerobic Roseobacters continue to carry denitrification genes (*29*) (table S1). As a first step towards bridging between our laboratory observations and the marine environment, we examined whether bacterial denitrification genes co-occur with high oxygen and chlorophyll levels in the ocean. Such co-occurrence could indicate the co-existence, and potential interaction, of phytoplankton and bacteria with NO-producing potential. We searched the Ocean Gene Atlas (OGA) (*50*), focusing on sampling points in the deep chlorophyll maximum layer (DCM) where photosynthesizing microalgae thrive, and specifically in locations where high levels of oxygen were measured (fig. S4A). We found that the genes *nirK* and *nirS*, both encoding nitrite reductases harbored by members of the Roseobacter group (*29*), are detected in these regions. In addition, genes encoding the NO reductase subunits, *norC* and *norB*, were found in all locations where *nirK* and *nirS* were detected (fig. S4B).

By examining the taxonomic distribution of bacteria in the DCM, we found that a high percentage of bacterial species that were detected in this oxygen-rich region and that carry denitrification genes, are from the *Rhodobacter* family, to which marine Roseobacters belong (fig. S4C). In addition, we identified metagenome-assembled genomes (MAGs) isolated from the ocean surface that belong to the Roseobacter group (*51*). A phylogenetic tree was built using the assembled MAGs, along with genomes of Roseobacters that were previously experimentally-validated to be either strictly aerobic (table S1, and references within) or facultatively anaerobic (table S3, and references within) (fig. S4D). Interestingly, the phylogenetic tree revealed that several MAGs grouped together with strict aerobes, and that these bacteria carry only part of the denitrification genes, but not all the genes essential for the full canonical denitrification pathway (Fig. 1A, fig. S4D). This is contrary to most anaerobes that do carry the entire enzymatic complement of denitrification genes. Taken together, the environmental data support a scenario in which bacteria that carry denitrification genes for reasons other than bioenergetics co-occur, and possibly interact, with photosynthesizing microorganisms. Therefore, bacterial NO production in oxygen-rich waters might be a common yet overlooked mechanism underlying interspecies interactions.

## Discussion

Our study demonstrates a novel route of microbial inorganic communication and highlights the need to expand our view beyond organic metabolic exchange. In our algal-bacterial system, a dynamic interaction begins with an initial mutualistic phase that transitions to a pathogenic phase as the algal culture ages (*28*). In the current study, we identify the algal nitrite signal that indicates algal growth phase and promotes bacterial transition to pathogenicity, we reveal and characterize the NO signal that allows bacteria to exert their pathogenicity, and we uncover how bacterial NO affects the algal population (Fig. 5).

**Fig. 5.**
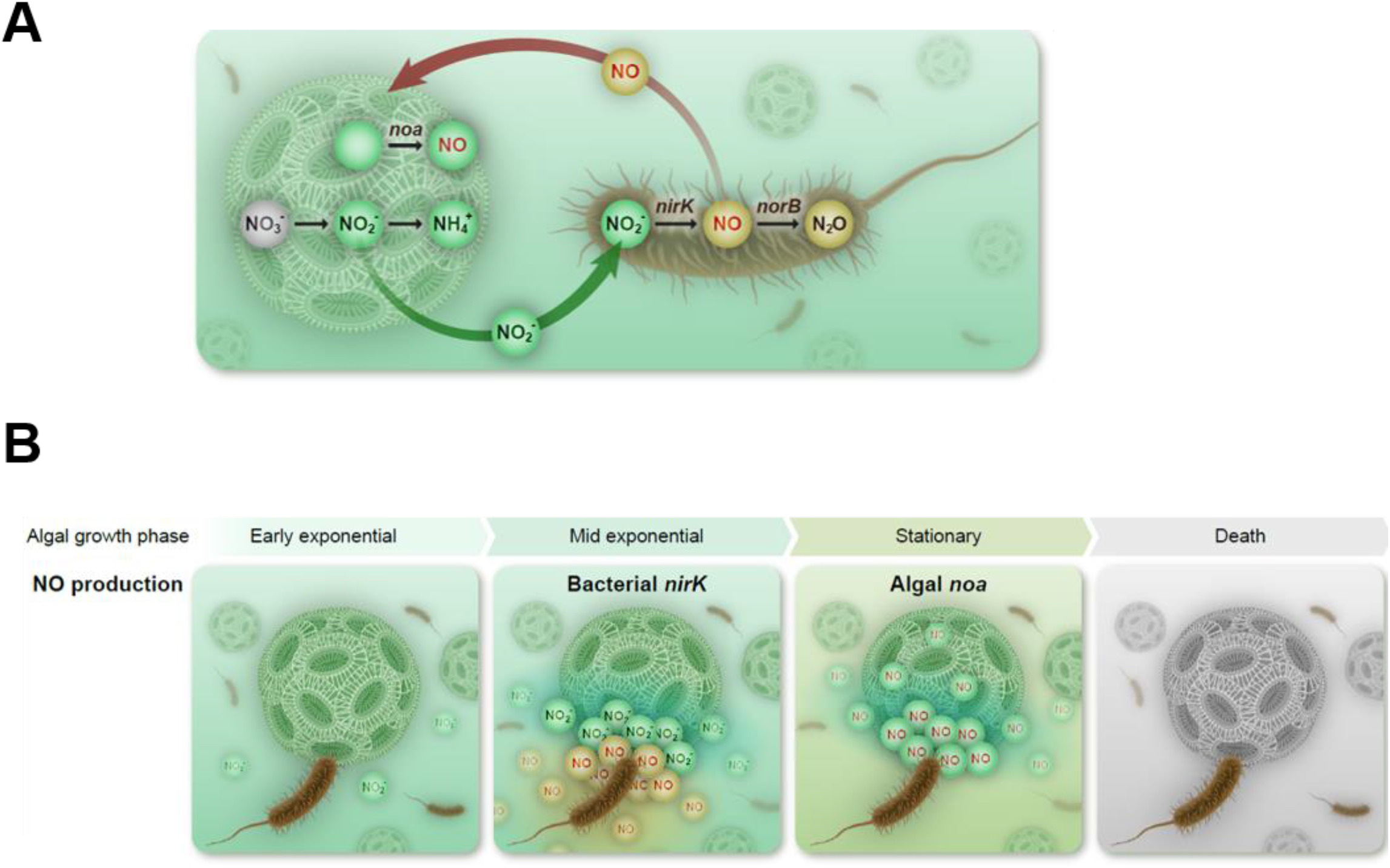
A model depicting inorganic nitrogen exchange during algal-bacterial interactions. **(A)** Microbial exchange of inorganic nitrogen. Algae secrete nitrite (NO_2_^−^) during assimilation of nitrate (NO_3_^−^) to ammonium (NH_4_^+^). Bacteria reduce nitrite to nitric oxide (NO) and possibly further to nitrous oxide (N_2_O), involving the expression of the denitrification genes *nirK* and *norB*, respectively. NO is secreted from bacteria and diffuses back to algae triggering algal NO production from an unknown precursor involving the expression of the *noa* gene. **(B)** Inorganic nitrogen exchange underlies the algal-bacterial dynamic interaction. Algae secrete nitrite in a growth-dependent manner, with a nitrite peak during mid exponential growth. Bacteria reduce algal-secreted nitrite to NO via denitrification involving expression of the *nirK* gene. NO secreted by bacteria triggers algal death at stationary phase, during which algae express *noa* genes and produce NO. Algal-secreted NO propagates the death signal among algal cells, causing the collapse of the algal population.

Our data demonstrate a novel role of denitrification genes in inter-species communication. In our algal-bacterial model system, the exchange of inorganic nitrogen species that are known denitrification intermediates, allows sensing and killing of a neighboring organism. Strictly aerobic *P. inhibens* bacteria harbor denitrification genes and express them under oxygen-rich conditions for reasons unrelated to bioenergetics (Figs. 2 and S2). The denitrification genes are unable to support growth of these bacteria under oxygen-depleted conditions (fig. S2C). Importantly, mutant bacteria that do not carry the denitrification genes are not capable of triggering death in the algal partner. Rather than being an ephemeral intermediate in respiration-related denitrification, NO appears to play a central role in the algal-bacterial interaction by triggering algal death.

Our results suggest a novel inorganic route in bacterial pathogenicity towards algae, independent of the previously described organic routes (*28, 35, 36, 44, 52-55*). Algae secrete nitrite during exponential growth, and bacteria reduce algal-secreted nitrite to NO which acts as the death agent that instigates the collapse of the algal population (Fig. 5). Our work highlights nitrite as an inorganic growth phase-specific signal, establishing a novel link between algal growth and induced bacterial pathogenicity. The various pathways described here and in previous studies (*28, 35, 36, 44, 52-55*) that enable bacteria to promote death of their algal partner point to the importance of this process in microbial interactions.

Our findings expand current knowledge on the role of NO in microbial ecology. Data from the current work and from previous studies demonstrate that NO is a signaling molecule in algae (*17, 43, 47, 56, 57*). We identified *noa* genes that likely encode NOA enzymes in *E. huxleyi*, one of which is up-regulated during algal death and potentially facilitates enhanced NO production (Fig. 4, fig. S3B). Algal NO appears to be central both in executing algal death (in a yet unknown molecular mechanism) and in propagating the death signal among the entire algal population (Fig. 5). Similarly, NO production by NOA enzymes in diatoms serves as a stress surveillance mechanism to alert the algal population (*47, 56*). Our observations shed new light on previous reports on the centrality of NO in the response of *E. huxleyi* to viral infection in non-axenic cultures (*17*). It is possible that in non-axenic cultures, bacterial NO derived from the leakage of nitrite from blooming algae triggers algal NO generation, which enhances the propagation of the viral infection in the algal population.

The role of NO as an extracellular signaling molecule is possible due to the close proximity between interacting algae and bacteria (*28*). Bacterial NO can diffuse into algal cells, efficiently acting as a signal that triggers algal death (*13*). In multicellular systems, local NO concentrations are crucial in determining specific intracellular responses of the affected cell (*58, 59*). Similarly, the differential expression of denitrification genes in bacteria is regulated by nanomolar changes in extracellular NO concentrations (*60*). In *E. huxleyi*, exposure to low NO concentrations increased population susceptibility to reactive oxygen species (*17*). Thus, the local concentration of NO in the algal phycosphere, the immediate volume surrounding the algal cell, might greatly influence algal fate and the nearby microbial population. Marine microbial “hot-spots”, such as marine snow particles and algal blooms, promote the development of local conditions with restricted diffusion, resulting in high concentrations of secreted compounds (*61-64*), which are otherwise diluted in the open ocean (*65*). Thus, short-lived, diffusible signals like NO might stimulate various localized responses, and are expected to differ between bloom and non-bloom conditions.

Data in our study indicate that aerobic bacteria can express denitrification genes for purposes other than respiration and energy production. This phenomenon appears of ecological importance beyond the empirical *E. huxleyi – P. inhibens* interaction. By mining the Ocean Gene Atlas (OGA) database, we were able to detect denitrification genes in oxygen-rich regions (fig. S4). The detected genes may belong to facultative anaerobes such as *P. inhibens* T5, a closely related species to our model bacterium, which harbors additional denitrification genes enabling anaerobic growth (table S3) (*66*). However, analysis of genomes of known strict aerobic bacteria, revealed that they harbor subsets of denitrification genes (table S1). Furthermore, expression of denitrification genes and the occurrence of active denitrification has been previously reported in aerobic conditions both in culture experiments and in environmental studies (*67-73*). The prevalence and ecological and biogeochemical significance of microbial denitrification in oxygen-rich environments remains to be determined. Furthermore, as microbial NO exchange can act across kingdoms, it might be an overlooked signaling molecule that shapes micro-scale to large-scale abundance and function of microbial populations across the global oceans.

## Acknowledgements

We are grateful for inspiring discussions with Prof. Roberto Kolter at the onset of this study. We thank Dr. Jörn Petersen for kindly providing the plasmid-cured mutant bacterial strain. We are grateful for the valuable input of Prof. Daniella Goldfarb on extracellular NO measurements by EPR. We thank Prof. Avigdor Scherz and Prof. Ayelet Erez for their input during early stages of this work. Finally, we are thankful to all members of the Segev lab for insightful comments and discussions.

## Funding

The Dean of Faculty fellowship (AA)

The Sir Charles Clore Postdoctoral Fellowship (AA)

Simons Foundation grant #622065 (IHZ, ARB)

The Israeli Science Foundation ISF 947/18 (ES)

The Peter and Patricia Gruber Foundation (ES)

The Minerva Foundation with funding from the Federal German Ministry for Education and Research (ES)

The Angel Faivovich Foundation for Ecological Research (ES)

The Weizmann SABRA - Yeda-Sela - WRC Program, the Estate of Emile Mimran, and the Maurice and Vivienne Wohl Biology Endowment (ES)

## Author Contributions

Conceptualization: AA, ARB, ES

Data curation: AA, MS

Formal analysis: AA, MS, RC, SBD, IHZ

Investigation: AA, MS, SBD, IHZ

Supervision: ES

Writing-original draft: AA, ES

Writing-review and editing: AA, ARB, ES

## Data and materials availability

All data are available in the main text or the supplementary materials.

