## Supplemental Materials and Methods, Figures S1-S4, Tables S1-S3. for "Aerobic Bacteria Produce Nitric Oxide via Denitrification and Trigger Algal Population Collapse"

### Strains and general growth conditions

The bacterial strain of *Phaeobacter inhibens* DSM 17395 was purchased from the German collection of microorganisms and cell cultures (DSMZ, Braunschweig, Germany). The bacterial Δ262 plasmid cured mutant was a kindly provided by the lab of Jörn Petersen, Leibniz Institut DSMZ, Germany (*42*). Bacteria were plated on ½ YTSS agar plates containing 2 g yeast extract, 1.25 g tryptone and 20 g sea salt per liter (all purchased from Sigma-Aldrich, St. Louis, MO, USA). Pure bacterial cultures were grown in CNPS medium consisting of L1-Si medium (see below) supplemented with glucose 5.5 mM, $Na_2SO_4$ 33 mM, $NH_4Cl$ 5 mM, $KH_2PO_4$ 2 mM, (all purchased from Sigma-Aldrich) *(28, 74)*. Cultures were incubated at 30°C shaking at 130 rpm.

The axenic algal strain of *Emiliania huxleyi* CCMP3266 was purchased from the National Center for Marine Algae and Microbiota (Bigelow Laboratory for Ocean Sciences, Maine, USA). Algae were grown in L1 medium according to Guillard and Hargraves (*75*), with the exception that $Na_2SiO_3$ was omitted following the cultivation recommendations for this algal strain, and the medium was referred to as L1-Si. Algae were grown in standing cultures in a growth room at 18°C under a light/dark cycle of 16/8 hr. Illumination intensity during the light period was 150 mmoles/m$^2$/s. Absence of bacteria in axenic algal cultures was monitored periodically both by plating on ½ YTSS plates and under the microscope.

Co-cultures of *E. huxleyi* and *P. inhibens* were cultured as follows: algal cell concentrations from a one week-old *E. huxleyi* culture was counted using a hemicytometer in a fluorescent microscope. An inoculum of $10^4$ algal cells was introduced into 30 ml of L1-Si medium and incubated as described above. After four days of algal growth, *P. inhibens* biomass from a plate was resuspended in 0.5 ml L1-Si, diluted X$10^4$ and 20 µl were added to 30 ml of algal culture, resulting in 10-100 colony forming units (CFU) per ml. The co-cultures were incubated in a growth room under the conditions described above for algal cultures. Sampling days are indicated as days following bacteria addition. When indicated, cultures were treated with the indicated concentration of the NO donor DEANO (Diethylammonium (Z)-1-(N,N-diethylamino)diazen-1-ium-1,2-diolate) diluted in ultrapure water (Cayman chemical, Ann Arbor, MI, USA). The NO scavenger c-PTIO (carboxy-PTIO) was added to a final concentration of 20 µM when indicated (ThermoFisher, Waltham, MA, USA).

**Monitoring algal growth in cultures**

Algal growth in cultures was monitored by a CellStream CS-100496 flow cytometer (Merck, Darmstadt, Germany), using 561 nm excitation and 702 nm emission. For each sample 50,000 events were recorded.

**Monitoring bacterial growth in co-cultures**

Bacterial growth in co-cultures was evaluated by sampling co-cultures at different time points, as indicated. Samples were serially diluted and plated on ½ YTSS plates. CFUs were counted and the concentration in the sampled culture was calculated. CFUs may include rosettes as well as individual bacteria.

**Monitoring bacterial growth in pure cultures**

Bacteria were grown in CNPS as described above. Cultures were monitored daily by $OD_{600}$ measurements in an Ultrospec 2100 pro spectrophotometer (Biochrom, Cambridge, UK) using plastic cuvettes. Cell numbers were calculated based on $OD_{600}$ values.

**Inorganic species measurements**

Axenic algal cultures were grown as previously described. On day of harvest, samples were filtered through a 0.1 µm syringe filter and analyzed using Thermo Scientific Gallery Plus discrete auto analyzer (ThermoFisher). Single atom concentrations were measured for each molecule from which Molar concentrations were calculated. L1-Si growth medium and filtered sea water were analyzed as controls. Standard ion concentrations were used for calibration.

**Extracellular nitrite measurements**

Extracellular nitrite was monitored using the Griess assay as follows; cultures were sampled on indicated days and filtered through a 0.2 µm syringe filter. Fresh standard curves were constructed using known concentration of $NaNO_2$ (Sigma-Aldrich). Samples and standards were diluted 1:1 using Griess reagent (Sigma-Aldrich), according to the protocol of the manufacturer, in a 96 well plate and incubated for 15 min in the dark. Samples were measured in triplicates at 545 nm using an Infinite M Plex plate reader (Tecan, Männedorf, Switzerland).

**Anaerobic growth**

*P. inhibens* and *E. coli* DH5α cells were plated on ½ YTSS agar plates supplemented with $NaNO_3$ or $NaNO_2$ at the indicated concentrations. Control plates were grown for 3 days at 30°C in aerobic conditions. Anaerobic growth was conducted by incubating plates for 1 month at 30°C, in a sealed BD GasPak™ container system, with BBL "GasPak" anaerobic pads, containing an oxygen depletion color indicator (BD, Franklin Lakes, NJ, USA).

**Quantitative real time PCR (qPCR)**

Bacteria were grown for 48 h in CNPS as detailed above. Cultures were treated with 10 µM nitrite for the indicated times prior to harvest. Pure algal cultures were grown for 48 h as indicated above. Cultures were treated with 100 µM of the NO-donor DEANO for 18 h treatment prior to harvest. Algal-bacterial co-cultures were grown as detailed above for the indicated times. For RNA extraction, $10^6$ algal cells or $10^8$ bacterial cells were harvested by centrifugation at 4000 rpm for 10 min. RNA was extracted using the Isolate II RNA mini kit (Meridian Bioscience, London, UK) according to the manufacturer instructions. Cells were ruptured in RLY buffer containing 1% β-mercapto-ethanol by bead beating with 100 µm low binding silica beads (SPEX, Metuchen, Netherland) for 5 min at 30 mHz. Approximately 1.4 µg of DNA was treated with 4µl Turbo DNAse (ThermoFisher), in a 50 µl reaction volume. RNA samples were cleaned and concentrated using RNA Clean & Concentrator™-5 kit (Zymo Research, Irvine, CA, USA) according to the manufacturer instructions. Algal-bacterial RNA samples were cleaned of algal ribosomal RNA using *E. huxleyi* riboPOOL™ kit (siTOOLs Biotech GmbH, Planegg, Germany) according to the manufacturer instructions, followed by additional RNA cleaning as described above.

Equal concentrations of RNA were utilized for cDNA synthesis using Superscript IV (ThermoFisher), according to manufacturer instructions.

qPCR was conducted in 384 well plates, using SensiFAST SYBR Lo-ROX Kit (Meridian Bioscience) in a the QuantStudio 5 (384-well plate) qPCR cycler (Applied Biosystems, Foster City, CA, USA). The qPCR program ran according to enzyme requirements for 40 cycles. Results were analyzed using a relative standard curve using the QuantStudio 5 software. Primer efficiencies were determined by qPCR amplification of serially-diluted cDNA. Only primer pairs with a minimum of 80% efficiency were selected, with the exception of Thioredoxin which was

65% efficiency. Samples were normalized using two housekeeping genes: bacterial housekeeping genes- *recA* and *gyrA* and algal genes for housekeeping proteins - *β-tubulin* and *rpl13*. DNA contamination was assessed by applying the same program on RNA samples that were not reverse transcribed. Relative gene expression levels were compared to non-treated samples grown under the same conditions. In co-cultures, relative gene expression levels were compared to expression levels on day 10.

Primer list of bacterial genes

| Gene name | Accession number | Forward Primer | Reverse Primer |
|---|---|---|---|
| *recA* | AFO91236.1 | GCTGACACCCAAGTCGGAG | AGCCGAACATAACGCCAATCT |
| *gyrA* | AFO91225.1 | GCCGATTCCTGACCTCCTTC | TCAGCTTATGTCGGGCTTCG |
| *nirK* | AFO93407.1 | CGGACAGTCAGATGGAACAC | ATTCATTCCGTGGGTGACAT |
| *norB* | AFO93403.1 | TGTTGACCGAGAAGTGGTGG | TTCCAGACCATCACGAAGGC |

Primer list of algal genes

| Annotated gene product | Accession number | Forward Primer | Reverse Primer |
|---|---|---|---|
| β tubulin | XM_005764044.1 | CAACATGAAGTGCGCCATCT | CCTCGGTGAACTCCATCTCG |
| Ribosomal protein L13 | XM_005781721.1 | ACCAGCACTTCCACAAGACG | TGCCGCAGCTTGTAGTTGTA |
| Glutathione synthase 3 | XM_005760150.1 | CTCCGGCAGGTCGAGCTAAA | GAGGCGTGCATGTCTTGCAG |
| L-ascorbate peroxidase | XM_005784352.1 | CGTGTCGACGCCTTAACAG | CACGATAGCCGGATGAGAAT |

| | | | |
|---|---|---|---|
| Autophagy protein 8 b | BK008761.1 | GGGCAGTTCGTGTACGTGAT | CTCGAGCTCTCCGAATGTGT |
| Thioredoxin | XM_005768338.1 | CACCAAGGCTGAGTTTGACA | GTAGAACTGGAAGGTCGGC |
| Metacaspase 3 | XM_005790343.1 | CCGACCACCAACTTCAACT | GGCTTTTCGTGGTAGTTGTC |
| Nitric oxide synthase 2 | MZ773650 | GTGGGTCTCGGTCGGATG | AGCCACCCCTGCTCCTAC |

**Light microscopy**

Fluorescence and phase contrast images were obtained using a Nikon Eclipse Ti2-E inverted microscope equipped with a CFI Plan Apochromat DM 100X objective lens (Nikon, Tokyo, Japan). All samples were spotted on thin 1% agarose pads for visualization at room temperature. Images were acquired using an Andor Zyla 4.2 camera controlled with Nis elements software. The 470/24 and 575/25 Lumencor Spectra X Chroma excitation filters were used. DAF-FM was captured using an ET519/26m filter. Images were processed identically for compared image sets.

**DAF-FM staining for microscopy**

Bacterial strains were grown for 24 h from an initial $OD_{600}$ of 0.04, in CNPS medium under the conditions described above. For the last 2 hours, bacteria were incubated with or without 100 µM $NaNO_2$. Cells were then centrifuged in 8000 rpm for 5 min at room temperature, resuspended in 10 µM DAF-FM Diacetate (4-Amino-5-Methylamino-2',7'-Difluorofluorescein Diacetate) (ThermoFisher) diluted in L1-Si, and incubated in the dark for 30 min. Cells were washed 3 times in L1-Si prior to imaging by light microscopy on agarose pads as described above. Image background was subtracted from images of bacterial cells which were not incubated with DAF-FM.

**Extracellular NO measurements**

Extracellular nitric oxide in bacterial and algal cultures was measured by Liposome-Encapsulated-Spin-Trap (LEST) and electron paramagnetic resonance (EPR) spectroscopy, based on the method previously published by Hirsh, *et al.* (*43*) with several adjustments. Multi Laminar Vesicles (MLVs) were prepared from a 9:1 molar ratio of the phospholipids POPC and DPPG (Avanti Polar

Lipids, Alabaster, AL, USA) in chloroform containing 2% methanol and 1% ultrapure water. The phospholipid mix was divided into glass bottles, evaporated under nitrogen flow and lyophilized overnight. Bottles were then capped and kept in -20°C for further use. LESTs were prepared one day prior to incubation with cells. On the day of preparation, the lipid film was dissolved in 1 ml buffer solution per 100 mg phospholipid of MLV Prep Buffer containing 10 mM MES pH 6.4 (Sigma-Aldrich) and 50 mM NaCl, 10 mM of the spin trap MGD (Santa Cruz Biotechnology, Heidelberg, Germany), and 2 mM ammonium iron (II) sulfate (Sigma-Aldrich). Glass bottles were capped with a rubber septum and the head space was flushed with nitrogen for 10 min. The lipid film was then completely suspended in the MLV Prep Buffer. Lipids were then frozen in liquid nitrogen and thawed in room temperature water five times to create frozen-and-thawed MLVs (FAT-MLVs). The rubber cap was then removed and EDTA at pH=6.4 (Sigma-Aldrich) was added to a final concentration of 2 mM. MLVs were then diluted 1:2 in NO Assay Buffer containing 20 mM HEPES at pH=7.4 (Sigma-Aldrich) and 140 mM NaCl, and centrifuged at 20,000 g for 30 min at 4°C. The supernatant was discarded and the MLVs were resuspended in NO Assay Buffer to a final volume of 2.5 ml. MLVs were loaded on a pre-equilibrated PD-10 de-salting column (GE Healthcare, Chicago, IL, USA) and collected with NO assay buffer in two 1.5 eppendorf tubes, followed by a second centrifugation at 20,000 g for 30 min at 4°C. MLVs were then resuspended at 480 µl volume per 100 mg phospholipid, and kept overnight at 4°C.

Axenic algal cultures or algal-bacterial co-cultures were grown as described above. Bacterial cells were grown for 24 hours from an initial $OD_{600}$ of 0.04, in CNPS medium under the conditions described above. On the day of measurement, bacterial and algal cultures were centrifuged at 4000 rpm and resuspended in 1 ml or 5 ml of growth medium, respectively. 30 µl of pre-made LESTs was added to each sample, as well as $NaNO_2$ when indicated. A sample treated with the NO-donor DEANO was used as a positive control for each experiment to verify proper LEST preparation and NO detection. Samples were incubated in the dark for 5 h. Then, samples were centrifuged at 20,000 g for 30 min at 4°C, resuspended in 100 µl, flash frozen in liquid nitrogen and kept at -80°C until measurement.

For EPR measurements samples were thawed and drawn up into microcapillary tubes. EPR spectra were recorded on a Bruker ELEXSYS E500 X-band spectrometer equipped with a Bruker ER4102ST resonator at room temperature. Experimental conditions were: 512 points, with microwave power of 20 mW, 0.1 mT modulation amplitude and 100 kHz modulation frequency.

Sweep range was 10 mT. NO concentrations were calculated according to a standard curve of known concentrations of Carboxy-proxyl. Background of an empty capillary was omitted from the measured values.

**DAF-FM measurements in algae**

1 ml of a one week-old algal culture was centrifuged at 8000 rpm for 5 min and stained with DAF-FM as detailed above. Cells were then washed with L1-Si and incubated for 30 min in the dark. Cells were washed again, resuspended in 400 µl of L1-Si and divided to aliquots of 50 µl. Each aliquot was diluted with 450 µl L1-Si as control or with 450 µl of bacteria, which were grown for 48h in CNPS as detailed above. Where indicated, bacteria were killed by treatment with gentamycin (100 µg/ml) and kanamycin (200 µg/ml) for 4 h. Bacterial death was validated by plating a sample of the antibiotic-treated bacteria on 1/2YTSS plates. 10 µM $NaNO_2$ was added where indicated. As a positive control for DAF-FM staining, stained algae were treated with 300 µM of the NO-donor DEANO. Samples were incubated in the dark for 2 h and then measured by flow cytometry. Detection of algal cells was conducted as described above. 10,000 algal events were collected. Detected algal events were plotted for DAF-FM fluorescence, using the 488nm excitation and 528 nm emission filter. Events of high DAF-FM fluorescence were gated according to the control of DEANO-treated algae. The relative percent of gated events was normalized to the background fluorescence of DAF-FM-stained algae with or without $NaNO_2$ treatment.

**RNA sequencing**

To identify PCD-related genes in *E. huxleyi* CCMP3266, we re-analyzed previously generated transcriptome data (*45*). In the previous work, we constructed a transcriptome of *E. huxleyi* CCMP3266 by integrating Illumina total RNA short-reads and PacBio full-length cDNA long-reads into a de novo assembled hybrid transcriptome (accession number: GIZZ00000000). In the same work, we collapsed redundant transcript isoforms into less redundant CCMP3266 gene loci, resulting in a "synthetic genome", which we used as reference for genetic analyses. In the current study, we used a text search to screen the *E. huxleyi* CCMP3266 transcript-based gene annotation table (Data S2 in (*45*)) for functions known to be hallmarks for PCD and oxidative stress. We further filtered the identified, putative PCD and oxidative stress genes for those that were differentially expressed (DE) during four different life phases of the alga in co-cultures with bacteria. The temporal DE analysis was conducted as previously described (*45*), using the Illumina total RNA sequencing data, and the "synthetic genome" of CCMP3266 as reference, but by applying a less stringent adjusted p-value cutoff of 0.1. The newly identified, putative *E. huxleyi* CCMP3266 PCD and oxidative stress genes, as well as their temporal expression during the transition from algal growth to demise, is presented as heatmap. The heatmap was generated with pheatmap (*76*), and rows were hierarchically clustered according to expression patterns with default options. Gene IDs and product annotations can be found in Data S2 in (*45*). Nucleotide sequences of gene transcripts can be retrieved from Data S1 of the same publication.

**Bioinformatical identification of algal *noa* genes**

The newly identified *noa* genes were predicted from *Emiliania huxleyi* genome sequence (*77*), and defined based on Illumina short read sequence (*78, 79*), as described in Feldmesser *et al* (*80*). Briefly, the sequences used for the initial searches were from *Phaeodactylum tricornitum* (a manually 5' extended version of Phatr3_J40200, the version 3 accession number of Phatr2_56150, and Phatr2_37004 or v3: Phatr3_EG00845.t1), Arabidopsis (NP_190329.2), and human (NP_000611.1, NP_000616.3, NP_000594.2). The protein sequences containing the YqeH domain, a conserved domain found in NO producing enzymes (*47*), were used for database searches (BlastP), multiple alignments (ClustalW 2.1) and phylogenetic trees (Neighbor-Joining in ClustalW, PhyML 3.0) (*81, 82*) to ensure their belonging to the correct family. *noa* sequences were aligned in MAFFT with the --leavegappyregion and --auto parameters (*83*). IQTree 1.6.12 was used to perform maximum-likelihood phylogenetic analyses with the best-fit model (LG+F+R5) according to the Bayesian information criterion and 100 bootstraps (*84*). The phylogenetic tree was visualized and annotated in FigTree, and branches were colored according to bootstrap values.

**Environmental data**

Environmental data was obtained using the Ocean Gene Atlas (OGA) server (*50*). The protein sequences of *nirK*, *norB* and *norC* from *P. inhibens* DSM 17395 and *nirS* from *Roseobacter denitrificans* were used as queries. Data analysis was limited to sampling points within the Deep Chlorophyll Maxima (DCM) region in which oxygen and chlorophyll measurements were available. Chlorophyll measurements were plotted against oxygen level measurements for each sampling point. Phylogenetic data was extracted from the taxonomic composition of each sampling point using Krona (*85*). The sampling point with the highest number of detected sequences (hits) in the DCM was chosen for data presentation. In the chosen sampling point, the numbers of hits for each gene was as follows: *nirK* 16, *norB* 50, *norC* 22.

**MAG and Roseobacter tree methods**

Metagenome-assembled genomes (MAGs) from the TARA Oceans Project were assembled in Delmont *et. al.* 2018 (*51*) and downloaded. MAGs grouping within the Rhodobacteraceae clade were retained for this analysis. Complete genomes from cultured anaerobes and aerobes in the Roseobacter clade were downloaded and both high-quality MAGs (over 70% complete and less than 10% contaminated) and cultured genomes were annotated using the PROKKA annotation software (*86*). Custom Hidden Markov Models for enzymes encoded by the phylogenetic gene *rpoB* or the denitrification genes *narG, nirS, nirK, norB,* and *nosZ* were searched against the annotated genomes to further verify annotations or find genes missed by the automated process in PROKKA. *rpoB* sequences belonging to each MAG or cultured genome were extracted and aligned in MAFFT with the L-INS-i method and --leavegappyregion parameter (*83*). IQTree 1.6.12 was used to perform maximum-likelihood phylogenetic analyses on the rpoB alignment with the best-fit model (LG+F+I+G4) according to the Bayesian information criterion and 100 bootstraps (*84*). The resulting tree was visualized in iTOL and annotated with the presence or absence of each denitrification gene within each genome.

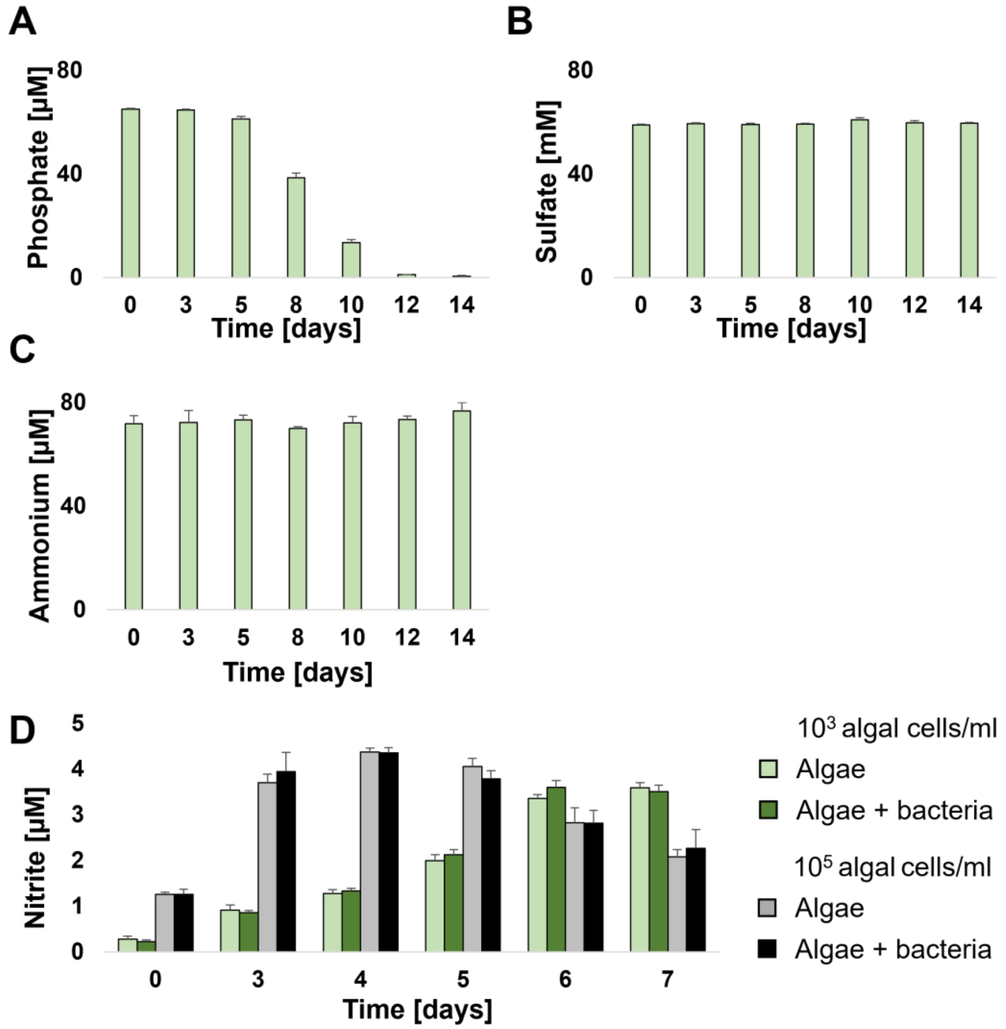


**fig. S1. Detection of inorganic molecules in filtrates of algal cultures.** (**A**) Phosphate (**B**) sulfate and (**C**) ammonium detected in filtrates of axenic algal cultures on the indicated days. (**D)** Nitrite levels measured in filtrates of axenic algal cultures and co-cultures inoculated with $10^3$ algal cells/ml (light and dark green bars) or $10^5$ algal cells/ml (gray and black bars). All results in the figure represent 3 biological replicates, error bars designate ± SD.

A

| Gene | Accession no. | Suggested activity |
|---|---|---|
| Nitrite reductase- *nirK* | WP_014881756.1 | Reduces nitrite to nitric oxide |
| Formate/nitrite transmembrane transporter | WP_014878953.1 | YfdC- Formate/nitrite transmembrane transporter |
| Putative nitrite/sulfite reductase | WP_014880398.1 | Sulfur metabolism |
| Nitrate/nitrite sensing domain containing protein | WP_014880942.1 | No suggested activity |

B

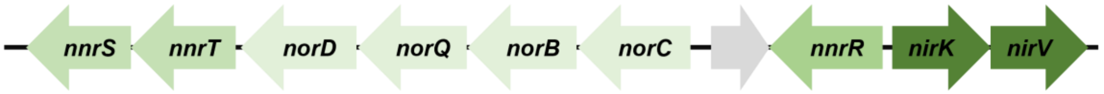


C

| **Strain** | ***Escherichia coli* DH5α** | | ***Phaeobacter inhibens* DSM 17395** | |
|---|---|---|---|---|
| **Growth conditions** | $+O_2$ | $-O_2$ | $+O_2$ | $-O_2$ |
| **½ YTSS** | + | + | + | - |
| **½ YTSS + 10mM $NaNO_3$** | + | + | + | - |
| **½ YTSS + 1M $NaNO_2$** | + | + | + | - |
| **½ YTSS + 500μM $NaNO_2$** | + | + | + | - |
| **½ YTSS + 100μM $NaNO_2$** | + | + | + | - |
| **½ YTSS + 20μM $NaNO_2$** | + | + | + | - |

D

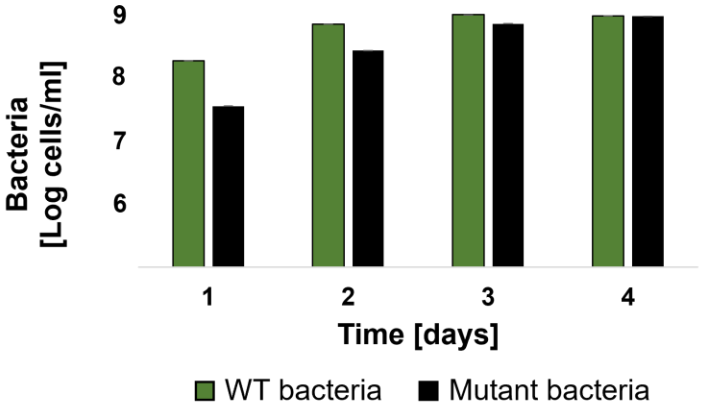

**fig. S2. Bacteria harbor denitrification genes but cannot grow under oxygen depleted conditions.** (**A**) List of genes, accession numbers and suggested activities related to nitrite metabolism in the *P. inhibens* genome. Data taken from the BioCyc database (*87*). (**B**) Denitrification-related genetic modules found in the 262 kb plasmid in the *P. inhibens* genome. Gray arrow- hypothetical protein. (**C**) Growth abilities of *P. inhibens* and *E. coli* on ½ YTSS plates supplemented with nitrogen species at the indicated concentrations. Plates were grown in oxygen-rich conditions (3 days) or oxygen-depleted condition (30 days). Growth indicated by (+). (**D**) Growth of WT (green bars) and mutant bacteria cured from the native 262 kb plasmid (black bars) in pure bacterial cultures. Each data point consists of 3 biological replicates, error bars designate ± SD.

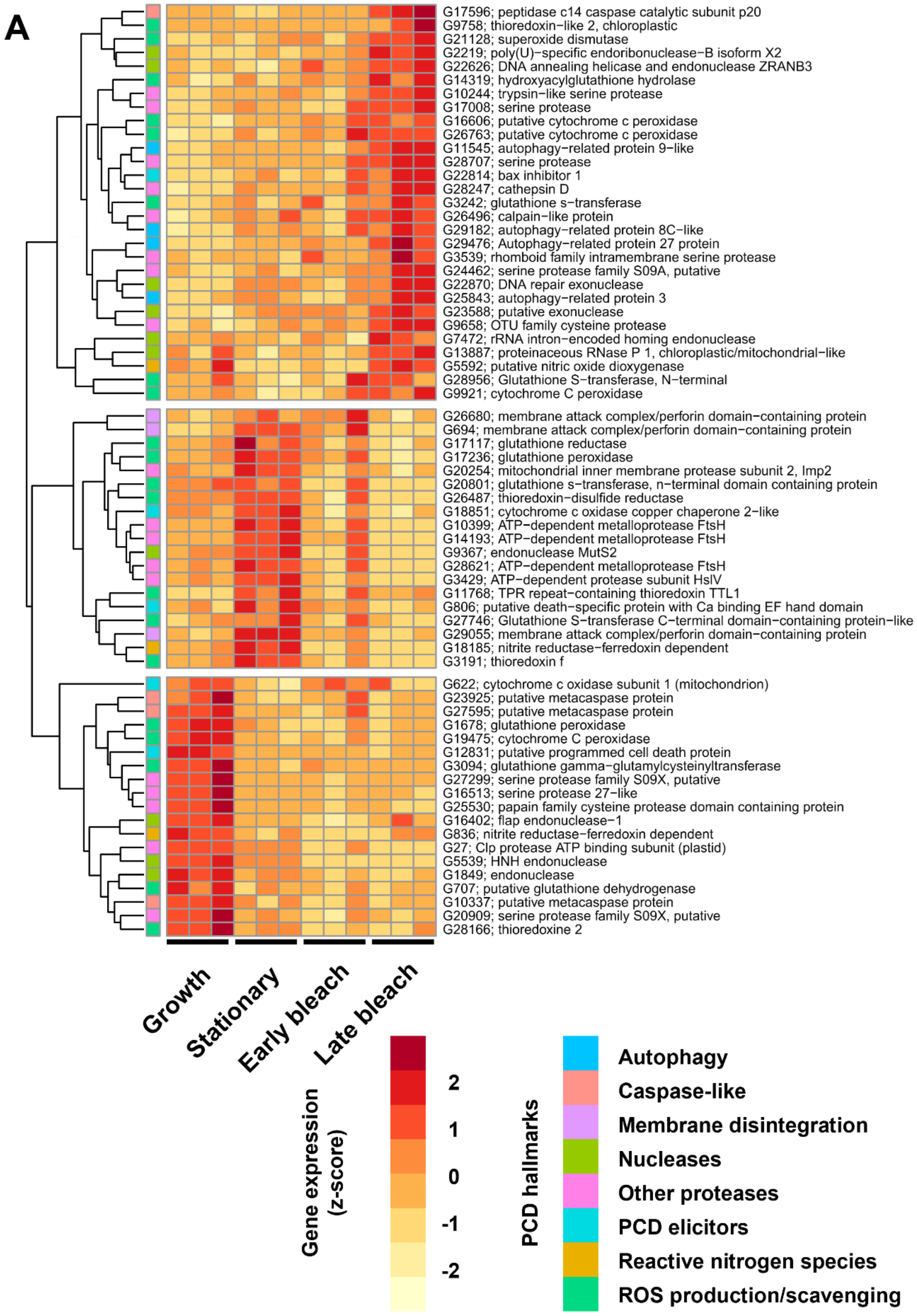
A
G17596; peptidase c14 caspase catalytic subunit p20
G9758; thioredoxin−like 2, chloroplastic
G21128; superoxide dismutase
G2219; poly(U)−specific endoribonuclease−B isoform X2
G22626; DNA annealing helicase and endonuclease ZRANB3
G14319; hydroxyacylglutathione hydrolase
G10244; trypsin−like serine protease
G17008; serine protease
G16606; putative cytochrome c peroxidase
G26763; putative cytochrome c peroxidase
G11545; autophagy−related protein 9−like
G28707; serine protease
G22814; bax inhibitor 1
G28247; cathepsin D
G3242; glutathione s−transferase
G26496; calpain−like protein
G29182; autophagy−related protein 8C−like
G29476; Autophagy−related protein 27 protein
G3539; rhomboid family intramembrane serine protease
G24462; serine protease family S09A, putative
G22870; DNA repair exonuclease
G25843; autophagy−related protein 3
G23588; putative exonuclease
G9658; OTU family cysteine protease
G7472; rRNA intron−encoded homing endonuclease
G13887; proteinaceous RNase P 1, chloroplastic/mitochondrial−like
G5592; putative nitric oxide dioxygenase
G28956; Glutathione S−transferase, N−terminal
G9921; cytochrome C peroxidase
G26680; membrane attack complex/perforin domain−containing protein
G694; membrane attack complex/perforin domain−containing protein
G17117; glutathione reductase
G17236; glutathione peroxidase
G20254; mitochondrial inner membrane protease subunit 2, Imp2
G20801; glutathione s−transferase, n−terminal domain containing protein
G26487; thioredoxin−disulfide reductase
G18851; cytochrome c oxidase copper chaperone 2−like
G10399; ATP−dependent metalloprotease FtsH
G14193; ATP−dependent metalloprotease FtsH
G9367; endonuclease MutS2
G28621; ATP−dependent metalloprotease FtsH
G3429; ATP−dependent protease subunit HslV
G11768; TPR repeat−containing thioredoxin TTL1
G806; putative death−specific protein with Ca binding EF hand domain
G27746; Glutathione S−transferase C−terminal domain−containing protein−like
G29055; membrane attack complex/perforin domain−containing protein
G18185; nitrite reductase−ferredoxin dependent
G3191; thioredoxin f
G622; cytochrome c oxidase subunit 1 (mitochondrion)
G23925; putative metacaspase protein
G27595; putative metacaspase protein
G1678; glutathione peroxidase
G19475; cytochrome C peroxidase
G12831; putative programmed cell death protein
G3094; glutathione gamma−glutamylcysteinyltransferase
G27299; serine protease family S09X, putative
G16513; serine protease 27−like
G25530; papain family cysteine protease domain containing protein
G16402; flap endonuclease−1
G836; nitrite reductase−ferredoxin dependent
G27; Clp protease ATP binding subunit (plastid)
G5539; HNH endonuclease
G1849; endonuclease
G707; putative glutathione dehydrogenase
G10337; putative metacaspase protein
G20909; serine protease family S09X, putative
G28166; thioredoxine 2
Growth
Stationary
Early bleach
Late bleach
Gene expression (z-score)
2
1
0
-1
-2
PCD hallmarks
Autophagy
Caspase-like
Membrane disintegration
Nucleases
Other proteases
PCD elicitors
Reactive nitrogen species
ROS production/scavenging

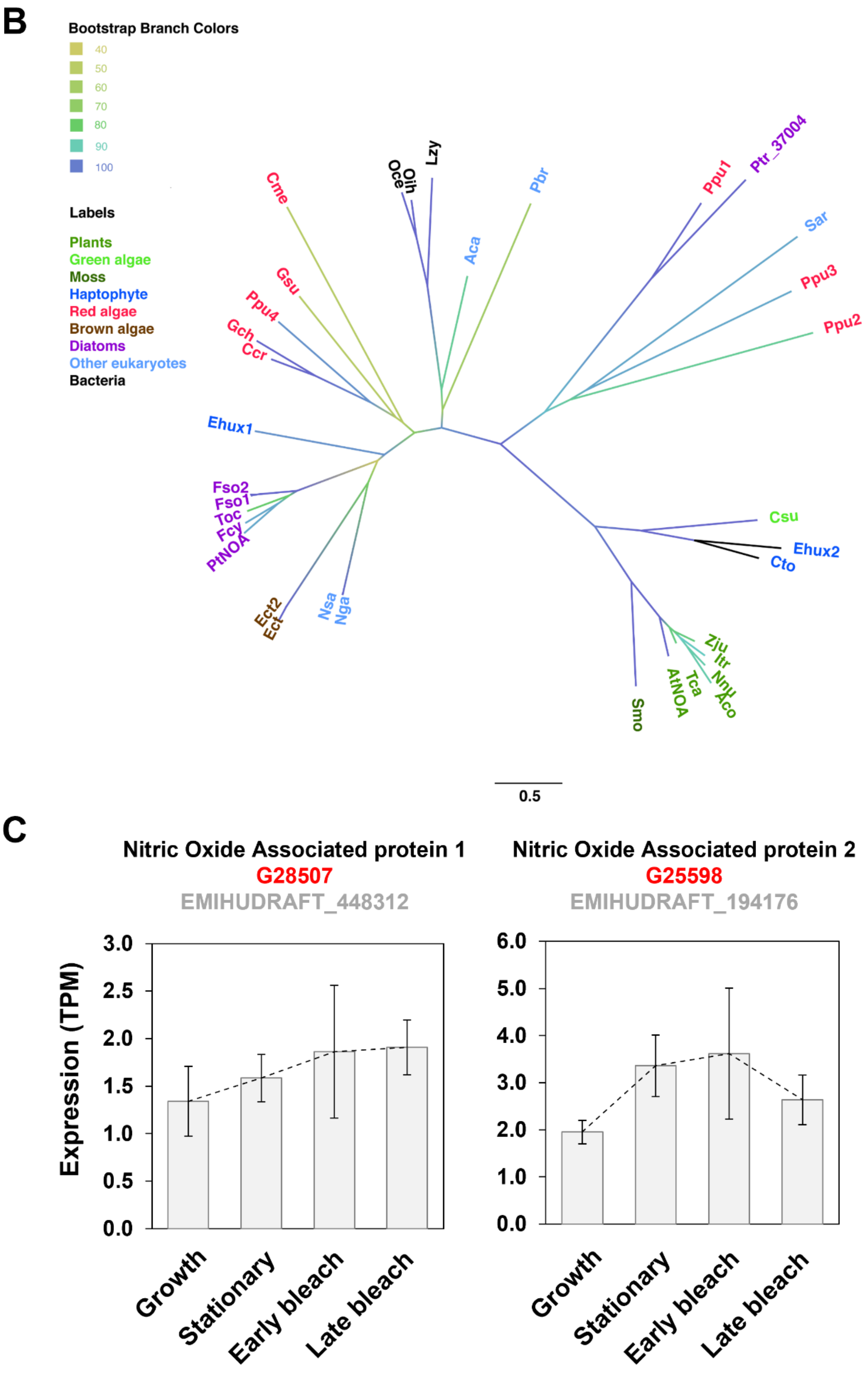

B
Bootstrap Branch Colors
40
50
60
70
80
90
100
Labels
Plants
Green algae
Moss
Haptophyte
Red algae
Brown algae
Diatoms
Other eukaryotes
Bacteria
Cme
Oce
Oih
Lzy
Pbr
Ppu1
Ptr_37004
Sar
Ppu3
Ppu2
Aca
Gsu
Ppu4
Gch
Ccr
Ehux1
Fso2
Fso1
Toc
Fcy
PtNOA
Ect2
Ect
Nsa
Nga
Csu
Ehux2
Cto
Smo
AtNOA
Tca
Zju
Itr
Nnu
Aco
0.5
C
Nitric Oxide Associated protein 1
G28507
EMIHUDRAFT_448312
Nitric Oxide Associated protein 2
G25598
EMIHUDRAFT_194176
Expression (TPM)
3.0
2.5
2.0
1.5
1.0
0.5
0.0
6.0
5.0
4.0
3.0
2.0
1.0
0.0
Growth
Stationary
Early bleach
Late bleach

**fig. S3. Identification of PCD-like and oxidative stress related genes in algae.** (**A**) A heatmap showing the expression of putative *E. huxleyi* oxidative stress and PCD-related genes that were differentially expressed during the indicated four phases of co-culture (adjusted p-value <0.1). Changes in gene expression are presented in red to yellow scale, with yellow indicating a decrease gene expression and red indicating an increase in gene expression. Genes were clustered according to expression pattern similarities. Gene IDs and product annotations are presented at the right side of the heatmap. Colored squares at the left side of the heatmap indicate suggested gene function. (**B**) Phylogenetic tree of Yqeh, the conserved domain in NO producing enzymes (see Materials and Methods). The branches are colored according to bootstrap, ranging from low (yellow) to high (blue). The species are colored according to phylogenetic range, see table S2. Plants- green, green algae- bright green, moss- dark green, haptophyte- blue, red algae- red, brown algae- brown, diatoms- purple, other eukaryotes- dark blue, bacteria- black. Tree scale: 1. (**C**) Gene expression levels of algal *noa* genes identified in (B) presented as transcripts per million (TPM), determined by total RNA sequencing on the indicated culturing phase. Gene annotations (bold), CCMP3266 gene IDs (red), and locus tag names of *E. huxleyi* CCMP1516 homologues genes present in the reference genome (gray). Expression data were taken from previously generated data (*45*). Each data point consists of 3 biological replicates, error bars designate ± SD.

**A**

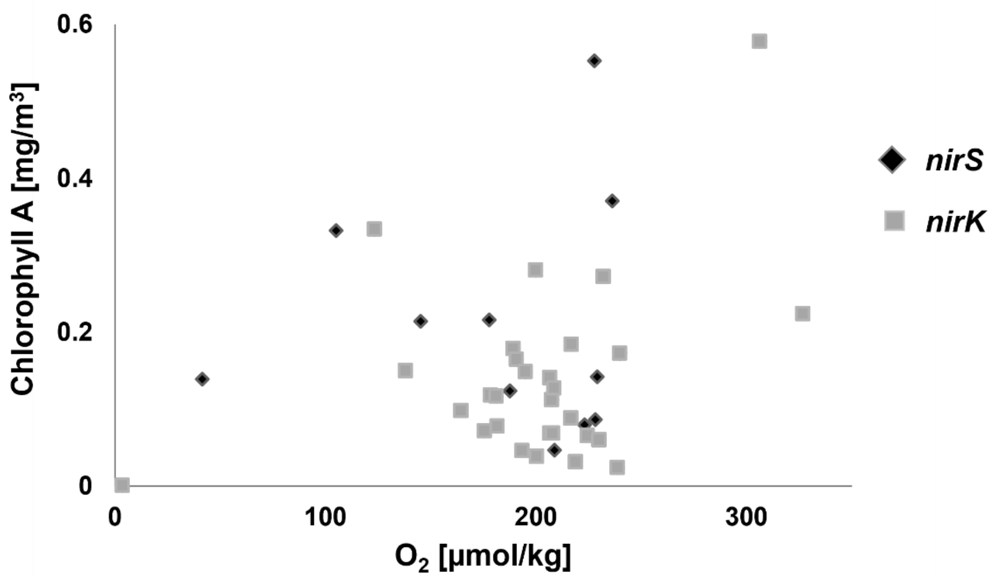


**B**

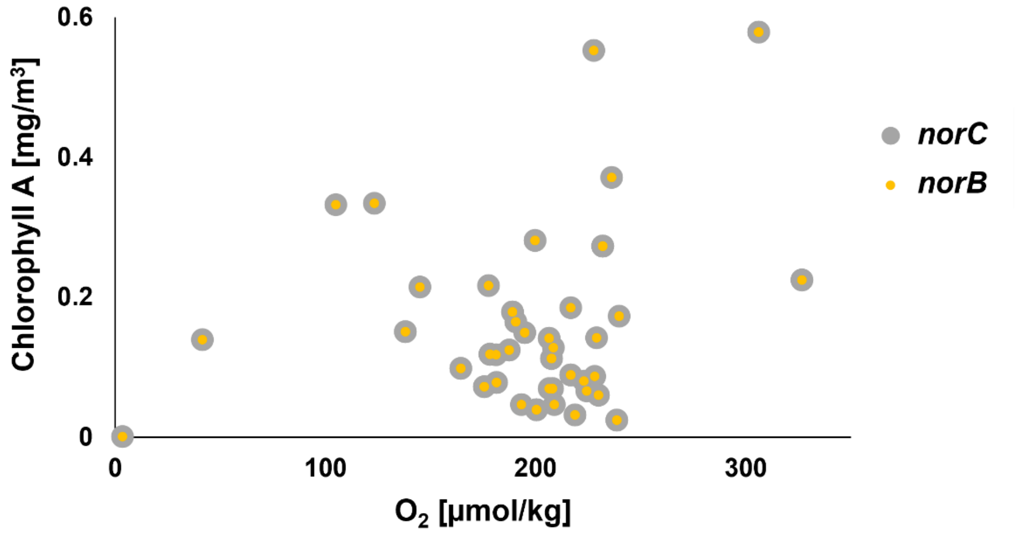


**C**

| Gene | *Rhodobacteriales* | *Rhodobacteracea* |
|---|---|---|
| *nirK* | 19% | 6% |
| *norC* | 14% | 9% |
| *norB* | 20% | 16% |

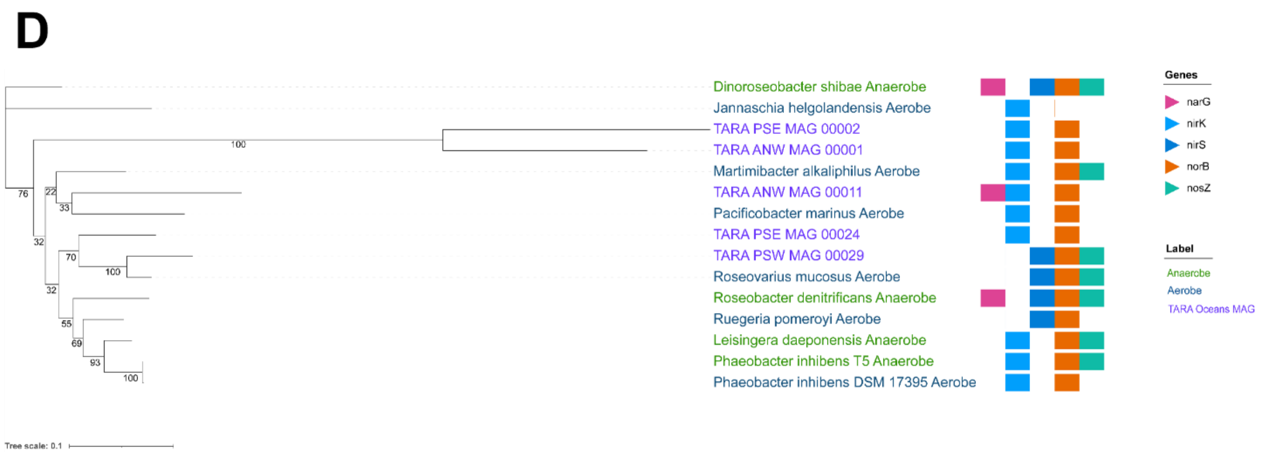


**fig. S4. Denitrification genes are found in deep chlorophyll maxima (DCM) regions in the ocean. (A-B)** DCM sampling points in which denitrification genes were detected are presented in dots, plotted against chlorophyll A and oxygen concentrations measured at the same location; (A) *nirS* and *nirK* (B) *norC and norB.* (**C**) Taxonomic distribution of the denitrification genes homologs detected in DCM sampling locations belonging to the *Rhodobacteriales* order and the *Rhodobacteracea* family. The location with the highest number of species detected was selected for presentation. (**D**) Phylogenetic tree of the *rpoB* gene from Roseobacter genomes of strict aerobes (table S1, species name in blue), anaerobes (table S3, species name in green) and Roseobacter MAGs (MAG identification name in purple). Bootstrap values are indicated at junctions. Identified denitrification genes in genomes are indicated in colorful squares to the right of each species. Pink- *narG* (nitrate reductase), light blue- *nirK* and dark blue- *nirS* (nitrite reductases), orange- *norB* (nitric oxide reductase), green- *nosZ* (nitrous oxide reductase). Tree scale: 0.1.

| Bacterial species | Location of isolation | Accession number | Reference |
|---|---|---|---|
| *Jannaschia helgolandensis* DSM 14858 | Surface water sample, North Sea, Germany | *nirK* WP_092763076.1 | (*88*) |
| *Roseovarius Mucosus* DSM 17069 | A dinoflagellate | *nirK* KGM86461.1 | (*89*) |
| *Martimibacter alkaliphilus* HTCC2654 | Western Sargasso Sea | *nirK* TYP84150.1 | (*90*) |
| *Pacificbacter marinus* CECT 7971 | Seawater sample, Yellow Sea, Korea | *nirK* WP_085849790.1 | (*91*) |
| *Rugeria pomeroyi* DSS-3 | Costal sea water | *nirS* AAV97354.1 | (*92*) |
| *Phaeobacter inhibens* DSM 17395 | Eukaryotic hosts and coastal biofilms(*93*) | *nirK* WP_014881756.1 | Current study |

**table S1. Strict aerobic bacteria that carry denitrification genes.** List of experimentally validated strict aerobic Roseobacters found in the literature, their location of isolation, identified denitrification gene and accession number, and relevant reference.

| Sequence name | Species | phylogenetic group | accession number | additional information |
|---|---|---|---|---|
| Ehux1 | *Emiliania huxleyi* | haptophyte | | |
| Ehux2 | *Emiliania huxleyi* | haptophyte | | |
| Aca | *Acanthamoeba castellanii str. Neff* | eukaryote | XP_004354073.1 | Ameobozoa |
| Aco | *Ananas comosus* | monocot | OAY76285.1 | pineapple |
| AtNOA | *Arabidopsis thaliana* | eudicot | NP_190329.2 | |
| Ccr | *Chondrus crispus* | red algae | XP_005713542.1 | |
| Cme | *Cyanidioschyzon merolae strain 10D* | red algae | XP_005535642.1 | |
| Csu | *Coccomyxa subellipsoidea C-169* | green algae | XP_005650839.1 | |
| Cto | *Chrysochromulina tobinii* | haptophyte | KOO31498.1 | |
| Ect1 | *Ectocarpus sp. CCAP 1310/34* | brown algae | CAB1112732.1 | |
| Ect2 | *Ectocarpus siliculosus* | brown algae | CBN74939.1 | |
| Fso1 | *Fistulifera solaris* | diatom | GAX26013.1 | |
| Fso2 | *Fistulifera solaris* | diatom | GAX24367.1 | |
| Gch | *Gracilariopsis chorda* | red algae | PXF48445.1 | |
| Gsu | *Galdieria sulphuraria* | red algae | XP_005706222.1 | |
| Itr | *Ipomoea triloba* | eudicot | XP_031117426.1 | trilobed morning glory |
| Lzy | *Lactobacillus zymae* | firmicute | WP_057734189.1 | Bacteria |
| Nga | *Nannochloropsis gaditana* | eukaryote | EWM29198.1 | Stramenopile |
| Nnu | *Nelumbo nucifera* | flowering plant | XP_010264327.1 | sacred lotus |
| Nsa | *Nannochloropsis salina CCMP1776* | eukaryote | TFJ88110.1 | Stramenopile |
| Oce | *Oceanobacillus sp. AG* | firmicute | WP_156858045.1 | Bacteria (fermented foods) |
| Oih | *Oceanobacillus iheyensis* | firmicute | WP_106896989.1 | Bacteria (deep-sea) |
| Pbr | *Plasmodiophora brassicae* | eukaryote | SPQ94216.1 | clubroot (plant pathogen, Rhizaria) |
| Ppu1 | *Porphyridium purpureum* | red algae | KAA8498098.1 | |
| Ppu2 | *Porphyridium purpureum* | red algae | KAA8499207.1 | |
| Ppu3 | *Porphyridium purpureum* | red algae | KAA8498134.1 | |
| Ppu4 | *Porphyridium purpureum* | red algae | KAA8490779.1 | |
| PtNOA | *Phaeodactylum tricornutum CCAP 1055/1* | diatom | XP_002184124.1 | |
| Ptr_37004 | *Phaeodactylum tricornutum* | diatom | Phatr3_EG00845.t1 Phatr2_37004 | |
| Sar | *Sphaeroforma arctica JP610* | opisthokont | XP_014153601.1 | |
| Smo | *Selaginella moellendorffii* | club-mosses | EFJ38034.1 | moss |
| Tca | *Theobroma cacao* | eudicot | XP_007038372.2 | cocoa bean |
| Toc | *Thalassiosira oceanica* | diatom | EJK65712.1 | |
| Zju | *Ziziphus jujuba* | eudicot | XP_015877800.1 | common jujube |

**table S2: List of species that carry a Yqeh domain**. List of species used to build the phylogenetic tree in fig. S3B. The name given to each species is color coded according to the phylogenetic range presented in fig. S3B. The phylogenetic group and gene accession number are also noted.

| Bacterial species | Isolated from | Reference |
|---|---|---|
| *Dinoroseobacter shibae* FL12 | A dinoflagellate | (*94*) |
| *Roseobacter denitrificans* | Seaweed | (*95*) |
| *Leisingera daeponensis* | Tidal flat sediment | (*96*) |
| *Phaeobacter inhibens* T5 | German Wadden Sea | (*66*) |

**table S3. Anaerobic bacteria.** List of anaerobic Roseobacters used for building the phylogenetic tree in fig. S4D, their location of isolation, and relevant reference.

**References** (74-96)
